# Antimicrobial activity of NK cells to *Trypanosoma cruzi* infected human primary Keratinocytes

**DOI:** 10.1101/2023.08.17.553656

**Authors:** Keshia Kroh, Jessica Barton, Helena Fehling, Hanna Lotter, Beate Volkmer, Rüdiger Greinert, Mouna Mhamdi-Ghodbani, Andrea Vanegas, Thomas Jacobs, Rosa Isela Gálvez

## Abstract

Infection with the protozoan parasite *Trypanosoma cruzi* is causative for Chagas disease, which is a highly neglected tropical disease prevalent in Latin America. Humans are primary infected through vectorial transmission by blood-sucking triatomine bugs. The parasite enters the human host through mucous membranes or small skin lesions. Since keratinocytes are the predominant cell type in the epidermis, they play a critical role in detecting disruptions in homeostasis and aiding in pathogen elimination by the immune system in the human skin as alternative antigen-presenting cells. Interestingly, keratinocytes also act as a reservoir for *T. cruzi*, as the skin has been identified as a major site of persistent infection in mice with chronic Chagas disease. Moreover, there are reports of the emergence of *T. cruzi* amastigote nests in the skin of immunocompromised individuals who are experiencing reactivation of Chagas disease. This observation implies that the skin may serve as a site for persistent parasite presence during chronic human infection too and underscores the significance of investigating the interactions between *T. cruzi* and skin cells. Consequently, the primary objective of this study was to establish and characterize the infection kinetics in human primary epidermal keratinocytes (hPEK). Our investigation focused on surface molecules that either facilitated or hindered the activation of natural killer (NK) cells, which play a crucial role in controlling the infection. To simulate the *in vivo* situation in humans, an autologous co-culture model was developed to examine the interactions between *T. cruzi* infected keratinocytes and NK cells. We evaluated the degranulation, cytokine production, and cytotoxicity of NK cells in response to the infected keratinocytes. We observed a strong activation of NK cells by infected keratinocytes, despite minimal alterations in the expression of activating or inhibitory ligands on NK cell receptors. However, stimulation with recombinant interferon-gamma (IFN-γ), a cytokine known to be present in significant quantities during chronic *T. cruzi* infections in the host, resulted in a substantial upregulation of these ligands on primary keratinocytes. Overall, our findings suggest the crucial role of NK cells in controlling acute *T. cruzi* infection in the upper layer of the skin and shed light on keratinocytes as potential initial targets of infection.

## Introduction

In Latin America, the infection caused by *Trypanosoma cruzi* (*T. cruzi*) and the subsequent development of Chagas disease is still the foremost zoonotic illness, having a top position among the most neglected tropical diseases recognized by the World Health Organization (WHO) (1, 2). This infection initiates within the epithelial layers of the skin and is accompanied by a transient local immune response until the parasites disseminate throughout the host’s body via the lymphatics or bloodstream. Towards the resolution of the acute infection, which typically spans two months, only a small number of parasites survive, concealed within specific target organs where they can persist for a lifetime. Cutting-edge bioluminescence techniques have recently identified the skin as a potential novel reservoir in mouse models. (3). Despite the crucial role of the skin as both the point of entry and a potential reservoir for *T. cruzi* parasites, there remains a lack of comprehensive characterization regarding the impact of the infection on the predominant cell types found within the skin, as well as the subsequent local immune responses. The skin’s complexity arises from the diverse array of somatic and immune cells, as well as the distinct functional layers it comprises. Therefore, analyzing these intricate interactions becomes imperative in order to comprehend the skin as a significant immunological organ (4, 5). The control of *T. cruzi* infection relies heavily on the pivotal role of NK cells and CD8^+^ T cells, as these immune cell types emerge as the central immune mediators in combating the infection, given the obligate intracellular nature of the parasite (6–9). This study aimed to characterize the impact of *T. cruzi* infection on human primary epidermal keratinocytes (hPEK) and assess the differential expression of immune regulatory ligands. First, to understand the kinetics of infection of hPEK, as well as the role of this cell type in orchestrating epidermal immune responses. Second, to gain insight into the specific mechanisms underlying the interplay between infected keratinocytes and the activation of NK cells. NK cells are a crucial immune cell population responsible for eliminating infected somatic cells in the skin and containing the parasite’s spread. Our findings demonstrate that *in vitro T. cruzi* infected hPEK can directly activate NK cells, resulting in increased secretion of pro-inflammatory cytokines (e.g., IFN-γ, IL-6) and cytolytic mediators (e.g., Granzyme A). Moreover, enhanced degranulation was observed through surface expression of CD107a. Utilizing the High Content Screening method developed in this study, we were able to quantify the increased cytotoxicity of NK cells. Additionally, our study shows that hPEK exhibited heightened expression of various HLA class I molecules and immune relevant ligands such as PD-L1, PD-L2 and ICAM-1 after infection and treatment with recombinant IFN-γ. Our study explores the relationship between hPEK and NK cells, uncovering the crucial role of keratinocytes in skin immunity against intracellular pathogens such as *T. cruzi*. The findings demonstrate that the direct interaction between infected keratinocytes and NK cells triggers cytokine production, cytolytic mediators, and enhanced cytotoxicity. This, in turn, may play a pivotal role in initiating a systemic robust immune response for effective pathogen defense.

## Results

### Human primary epidermal keratinocytes are an appropriate model for investigating the dynamics of *T. cruzi* infection within the skin

First, we aimed to study the kinetic of the *in vitro* infection of hPEK with two different strains of *T.cruzi*: Tulahuen and Brazil. The different infection stages were evaluated first by immunofluorescence staining 24 to 96 h post infection (p.i.) (**Fig. 1**). While at 24 h p.i. single intracellular *T. cruzi* amastigotes were visible, the number of intracellular amastigotes per cell increased over time. The timepoint 72 h p.i. was chosen for subsequent infection experiments, as the amastigotes have filled up the cells but were not yet transforming into the blood trypomastigote stage, as seen 96 h p.i.. The infection kinetics of both *T. cruzi* strains were similar, although the infection rates for *T. cruzi* Tulahuen were lower **(S1)**. To confirm and automatically quantify infection rates the Opera Phenix® HCS system was used and an image analysis sequence with the Harmony® software was established as described in previous study by Fehling et al. (10). To assess the infection rates at different multiplicity of infection (MOI), keratinocytes were seeded and infected with MOIs of 1:1, 3:1 and 6:1, and an infection period of 72 h. Cells were fixed and stained with DAPI (blue), and *T. cruzi* parasites were stained using polyclonal mouse anti-*T. cruzi* serum and a conjugated anti-mouse secondary antibody (red)). The anti-*T. cruzi* antibody also displayed a weak, unspecific staining of cells, which was used to identify the cytoplasm. The image analysis sequence was shown to robustly detect the cells and intracellular trypanosomes (**Fig. 2 A, B**). *T. cruzi* Brazil showed an infection rate of 6 % at a MOI of 1:1. This was increased to 17 % and 25 % with MOIs of 3:1 and 6:1, respectively. The infection rates of *T. cruzi* Tulahuen were much lower and were 1 % at an MOI of 1:1, 2 % at an MOI of 3:1, and 3 % at an MOI of 6:1. The average number of trypanosomes per infected cell was also higher in *T. cruzi* Brazil infected cells and ranged from 2.5 to 3.6. while the average number of *T. cruzi* Tulahuen trypanosomes was within the range of 1.3 to 2.1 (**Fig. 2C**). These results show that hPEK can be productively infected with *T. cruzi,* independently of the strain and allow further characterization regarding the interaction with NK cells. In the following experiments only the *T. cruzi Brazil* strain was used, since it led to more efficient infection.

**Figure 1.**
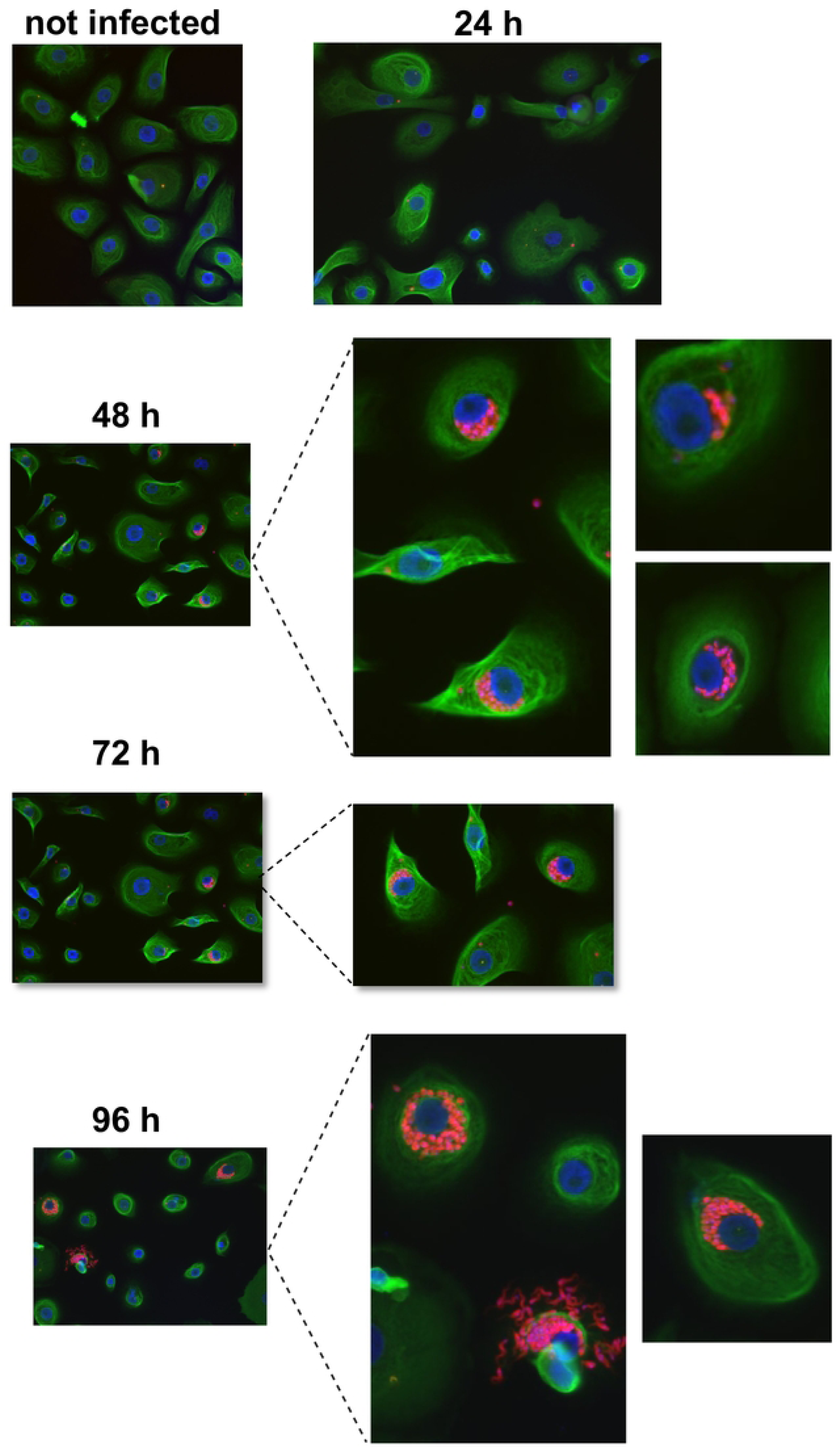
Infection of human primary keratinocytes with *T. cruzl* Brazil. Primary keratinocytes were infected with *T. cruzi* Brazil at a MOI of 3:1 for 24 h, 48 h, 72 h, and 96 h. Keratinocytes (green) and trypanosomes (red) were visualized by indirect immunofluorescenceusing a pan anti-cytokeratin antibody, polyclonal anti-T. *cruzi* serum, and DAPI. Images were obtained at 200x magnification. n.i., not infected; TcB inf., *T. cruzi* Brazil infected.

**Figure 2.**
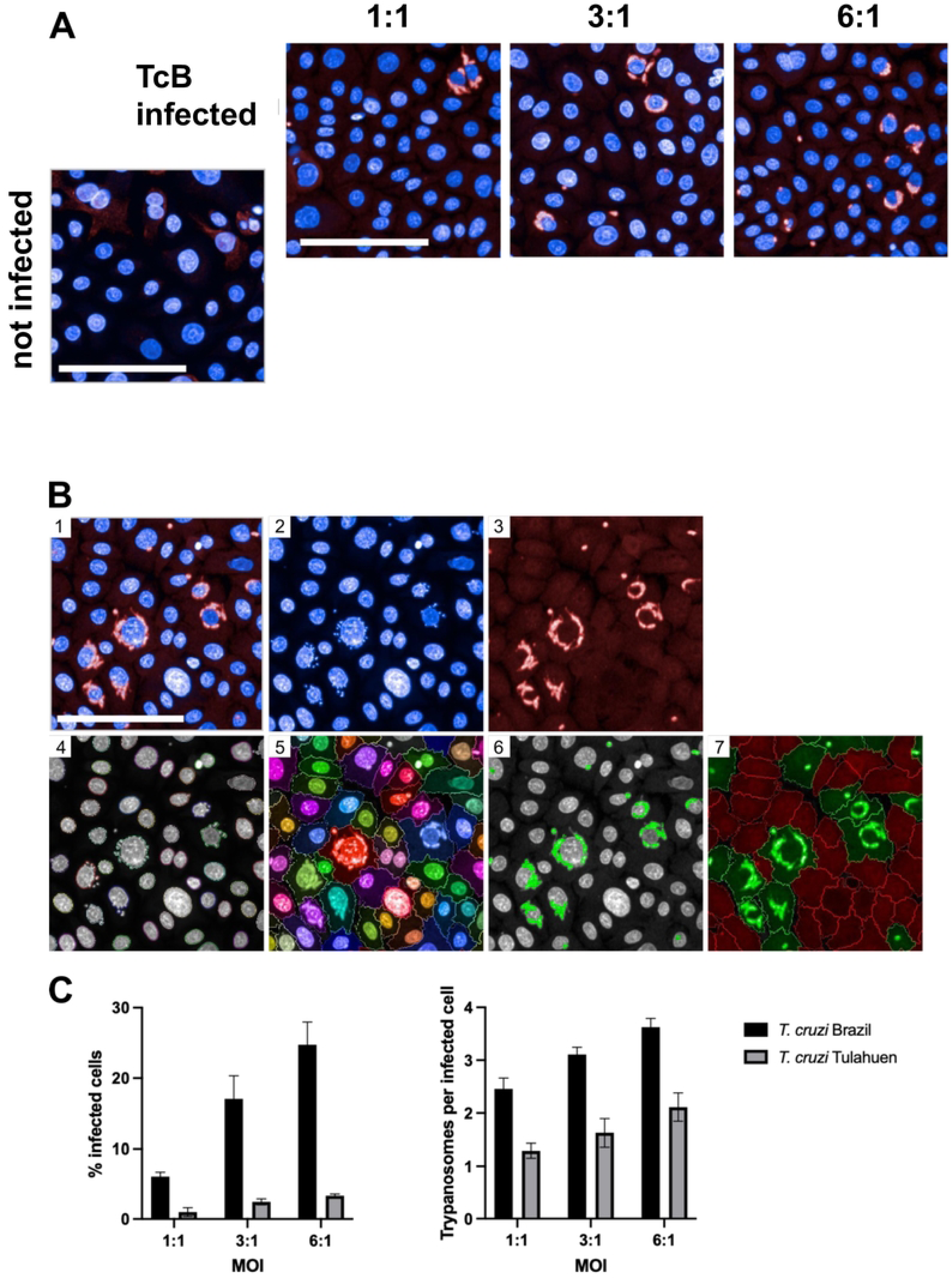
Evaluation of infection per cell usingthe high-<:ontent screening system Opera Phenix•. Primary keratinocytes were used to determine the infection rates of *T. cruzi* Brazil and *T. cruzi* Tulahuen 72 h p.i. at an MOI of 1:1, 3:1, and 6:1. *T.cruzi* parasites were stained using a murine polyclonal anti-T. *cruzi* serum and an AF647-conjugated anti-mouse lgG secondary antibody (red). Nuclei were stained with DAPI (blue). Images were obtained using the Opera Phenix" confocal imaging system. A) Representative image sections showing primary keratinocytes infected with *T. cruzi* Brazil, MOI of 1:1, 3:1, and 6:1. B) Image analysis sequence for one representativeimage*(T. cruzi* Brazil infected, MOl 6:1). (1) Input image, (2) nuclei staining (DAPI, 405 nm) (3) anti-T. *cruzi* staining (AF647, 640 nm), (4) detection of nuclei and (5) cytoplasm, (6) detection of trypanosomes, (7) selection of infected cells (green) and uninfected cells (red). C) Infection rates and average number of trypanosomes per infected cell for *T.cruzi* Braziland *T.*cruziTulahuen infected primary keratinocytes.5 wells were analyzed for each condition, and values are shown as mean± SD. Scale lOOµm.

### Autologous co-cultivation of *T. cruzi* infected hPEK activates NK cells and triggers cytotoxic activity

As keratinocytes may be one of the first infected cell types in the acute *T. cruzi* infection, an autologous model was employed to investigate the direct interaction between *T. cruzi* infected hPEK and NK cells from the same donor. A scheme of the co-culture experiments is shown in **(S2)**. After 24 hours, supernatant of the co-culture was collected for analysis of soluble mediators and NK cells were collected for flow cytometric analysis. The gating strategy and representative dot plots are depicted in **(S3)**. CD107a, which is exposed on the cell surface as a result of the degranulation of NK cells, serves as a marker for cytotoxicity (11). The background expression of CD107a by NK cells without target cells was subtracted to evaluate the fraction of CD107a^+^ NK cells. Using this method, an average of 7.5 % of NK cells co-cultured with non-infected keratinocytes, were positive for CD107a. In contrast, the proportion of CD107a^+^ NK cells in co-culture with infected keratinocytes increased to 24.1 %, exhibiting a threefold increase compared to the control. In addition, CD16 was analyzed since diminished CD16 expression can occur following activation of NK cells (12). CD16 was significantly downregulated on NK cells co-cultured with infected keratinocytes compared to non-infected keratinocytes (78.5 % vs. 88.3 % of CD16^+^ NK cells) (**Fig. 3 A**). A key protective mechanism employed by NK cells involves the release of cytokines and cytolytic mediators, such as lytic granules. To examine this, a multiplex bead-based immunoassay was conducted on the co-culture supernatants. The fold change in protein concentration was determined by comparing the mean of non-infected controls for each donor and experiment. IL-6, IFN-γ, and Granzyme A (GzmA) exhibited statistically significant differences (**Fig. 3 B**). Notably, the average fold change was higher in the supernatants of NK cells co-cultured with infected keratinocytes for all assessed cytokines and cytolytic mediators **(S4)**. Overall, the observed significant increase in activation and frequency of degranulated CD3^-^ CD56^+^CD107a^+^NK cells upon co-culturing with *T. cruzi* infected hPEK strongly supports the notion that infected hPEK can effectively stimulate secretion of pro-inflammatory cytokines and cytolytic mediators by NK cells.

**Figure 3.**
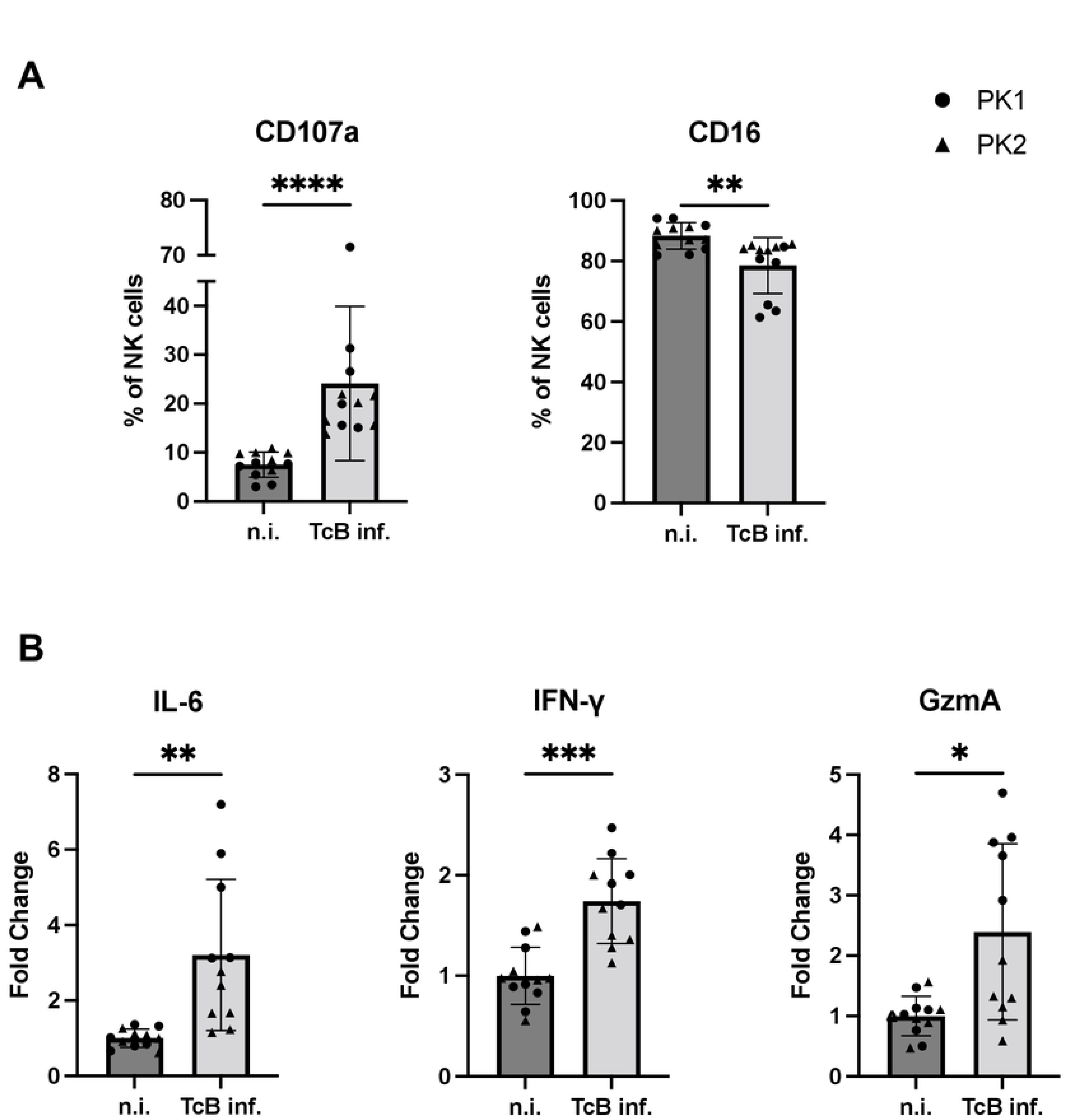
Activation of NI<cells co-cultured with *T. cruzi* Brazil infected keratinocytes. A) Frequency of CD107a• NK cells (left) and frequency of CD16• NK cells (right) after an autologous co­ culture with non-infected (n.i.) or *T. cruzi* Brazil infected (TcB) keratinocytes. B) Release of cytokines and cytolytic mediators by the NK cells.The fold change was calculated in relation to the non-infected controls. n = 11-12 from 2 donors (PKl, PK2) and 2 experiments each. Data are shown as mean± SD. Welch’s t tests were performed for normally distributed values, while Mann-Whitney tests were performed for values that did not follow normal distribution. * p s 0.05; •• p s 0.01; ••• p s 0.001; •••• p$ 0.0001; n.i. - non­ infected, TcB inf. - *T. cruzi* Brazil infected.

### High-content screening assay confirms killing of infected hPEK by NK cells

The cytotoxic effects of NK cells were assessed using the Opera Phenix® high-content screening system. Confocal images of hPEK were captured after co-culturing, and a cell counting analysis sequence was employed. The evaluation included determining the percentage of infected cells and the number of trypanosomes per infected cell. Diverse effector-to-target ratios (E:T ratios), ranging from 1:10 to 10:1 were analyzed. For this, the analysis sequence was modified to exclude NK cells, which were identifiable via the much smaller nucleus compared to the nuclei of hPEKs. Image sections of infected and non-infected keratinocytes from one donor, co-cultured without NK cells or with NK cells at E:T ratios of 1:1 and 10:1 is depicted in **Fig. 4 A**. For both donors, a significant reduction in the number of trypanosomes per infected cell was observed at the highest E:T ratio of 10:1. A killing effect of NK cells on intracellular trypanosomes was observed, as both the percentage of infected cells and the number of trypanosomes per infected cell were significantly decreased with the addition of NK cells (**Fig. 4 B**).

**Figure 4.**
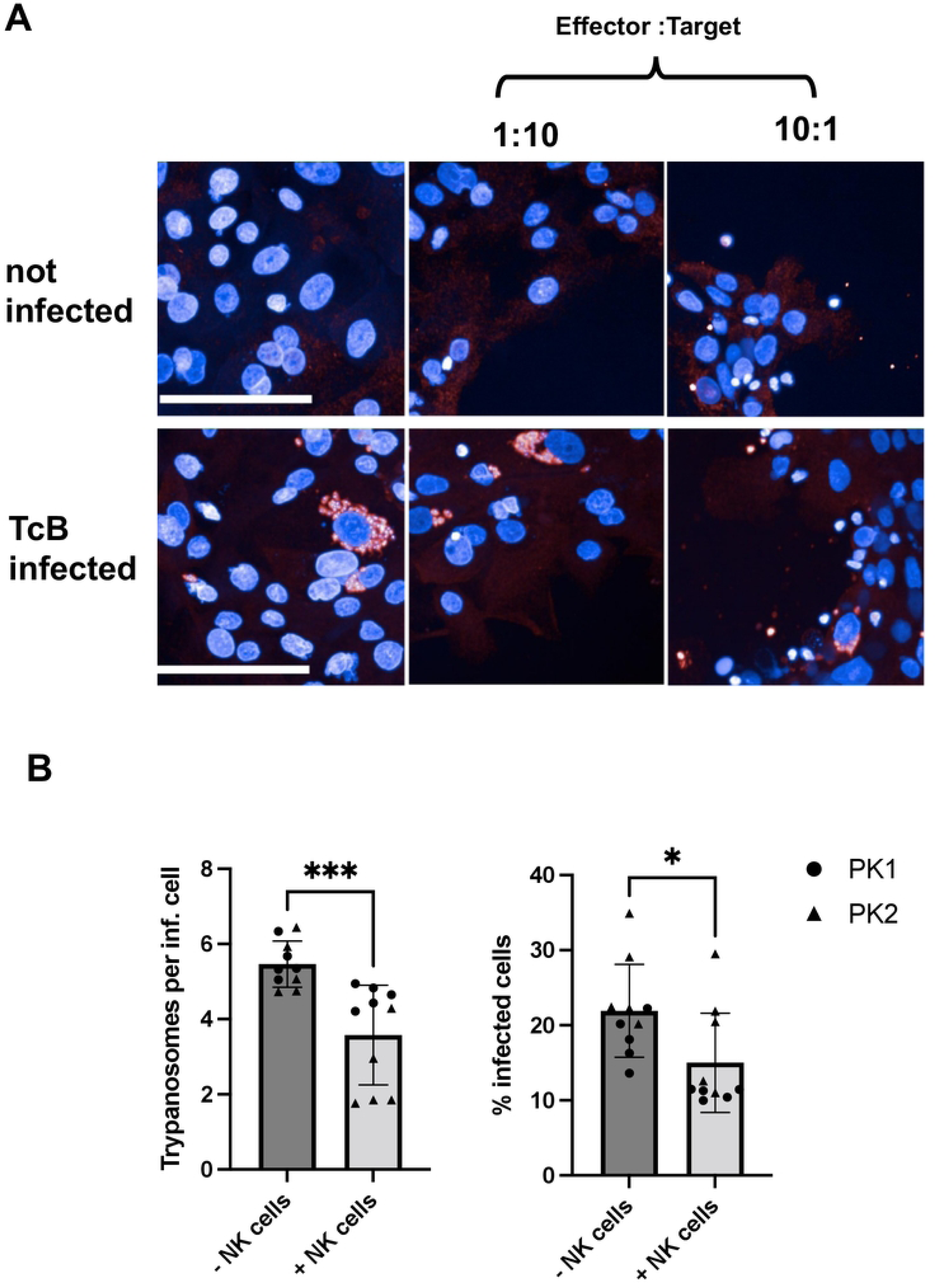
Trypanocidal effect of NI< cells on *T.*cruz/-infectedkeratlnocytes. A) Representative image sections of infected and non-infected keratinocytes co-cultured with NK cells at E:T ratios of 1:1 and10:l. B) Killing effect after co-culture with NK cells is depicted as percentage of infected cells and number of trypanosomes per infected cell. This was calculated using the Opera Phenix" HCS system. NK cells were added at an E:T ratio of 10:1.n = 10 from 2 donors (PKl, PK2), scale bar 100 µm. Data are shown as mean± SD. Mann-Whitney tests were performed to calculate statistical significance. • p $ 0.05; ••• p s 0.001.

### Upregulation of immune modulatory molecules in T*. cruzi* infected and IFN-γ stimulated human primary keratinocytes

To determine the surface expression of various molecules in hPEK infected with *T. cruzi and* stimulated with IFN-γ, flow cytometry analysis was conducted. The experimental design is shown in **(S5)**. The hPEK were infected at a MOI of 3:1 or stimulated with 0.1 ng/ml and 10 ng/ml of IFN-γ respectively. IFN-γ was chosen due to its crucial role in *T. cruzi* infection in murine models (13) and also chronic Chagas patients (14). We focused on the analysis of activating and inhibitory ligands as well as in HLA-molecules that mediate the interaction with NK cells. The gating strategy and representative dot plots are depicted in **(S6 A, B** for activating ligands, **C** for inhibitory ligands, **D** for HLA-Molecules).

The determination of the percentage of cells expressing the ligand was established using the negative control cell population. Consequently, the values should be interpreted as the percentage of cells expressing the target minus the background expression observed in untreated cells. Stimulation with 10 ng/ml of IFN-γ significantly increased the surface expression of nearly all analyzed targets, while the effects of *T. cruzi* infection or stimulation with 0.1 ng/ml IFN-γ were less pronounced. Specifically, the fraction of hPEK expressing the activating ligands HLA-DR and ICAM-1, and the inhibitory ligands PD-L1, PD-L2, and HVEM approached 100% after stimulation with 10 ng/ml of IFN-γ (**Fig. 5**).

**Figure 5.**
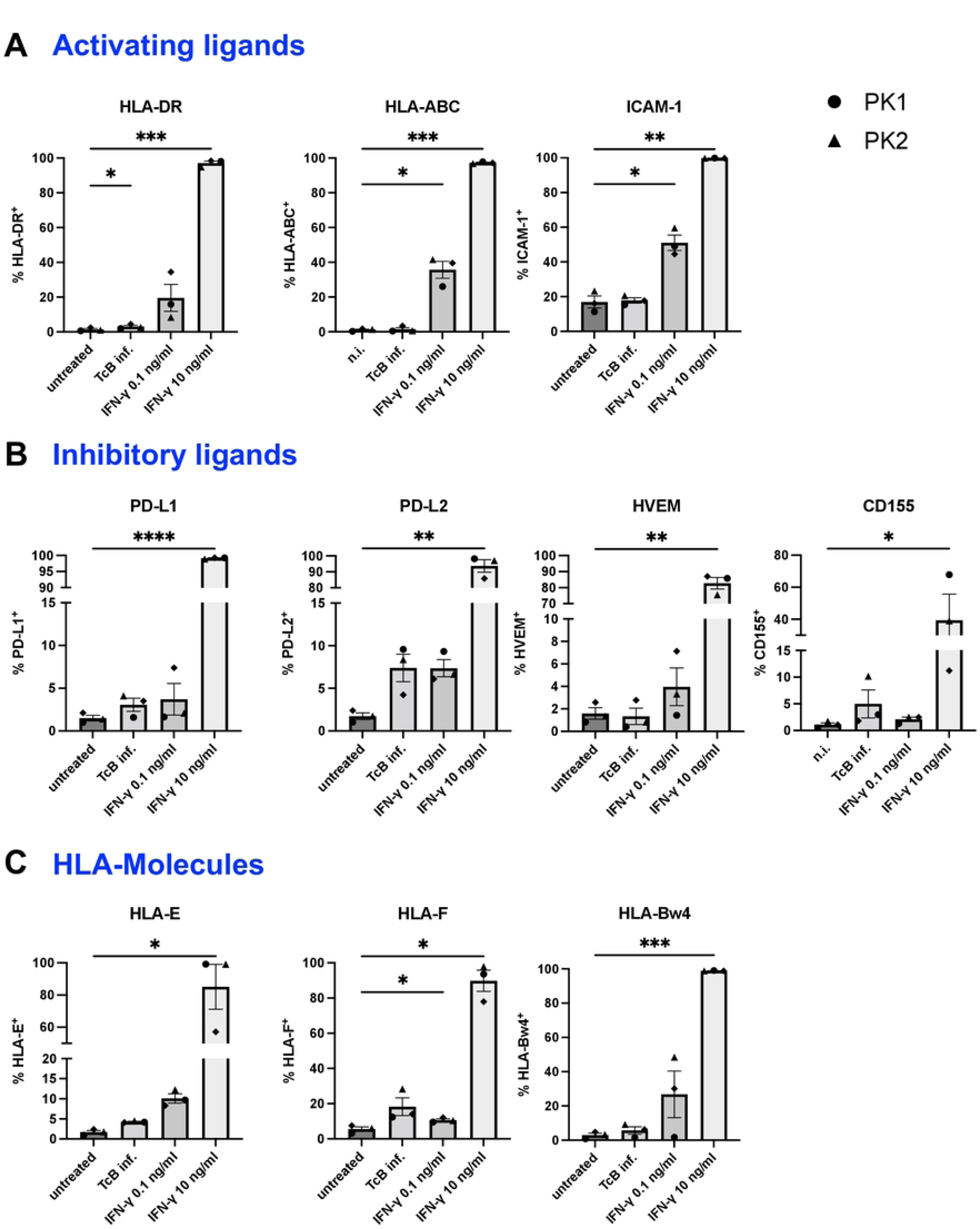
Immune modulatory molecules expression on infected and stimulated hPEI. HPEK were infected with *T. cruzi* Brazil at an MOI of 3:1 or stimulated with 0.1 ng/ml or 10 ng/ml of IFN-v in duplicates. Untreated keratinocytes were used as control. 72 h after infection/stimulation, the keratinocytes were detached and stained for flow cytometric analysis. A) Frequency of keratinocytes expressing the different activating ligands, B) Inhibitory ligands, C) HLA-Molecules. Statistical differences of untreated vs.infected/stimulatedcells were analyzed using Welch’s t tests for normally distributed values, and Mann-Whitney tests for values that did not follow normal distribution. n=3 from three donors. Data are shown as mean± SD. * p $ 0.05; •• p s 0.01; ••• p $ 0.001; •••• p$ 0.0001. TcB inf. - T. cruzi Brazil infected. Supplementary Figures

The fraction of cells expressing MIC-A/B and Galectin-9 was also higher compared to untreated controls but did not reach statistically significance **(S6 B, C)**. However, statistical significance might not be achieved due to high variation between donors. With the exception of CD155, increased expression was also observed for all analyzed targets when cells were stimulated with a lower concentration of IFN-γ (0.1 ng/ml). This increase was statistically significant for HLA-ABC, HLA-DR, PD-L2, and ICAM-1. The fraction of cells positive for HLA-DR and ICAM-1 was 19.6% and 51%, respectively, while for the other targets, it ranged from 3.7% to 7.4%. The T. *cruzi* infection had a lesser impact compared to IFN-γ stimulation, although significant induction of PD-L1, PD-L2, and MIC-A/B on infected hPEK were observed. Taken together, both the upregulation of surface molecules with activating and inhibitory effects was observed. The IFN-γ stimulation induced significant upregulation of almost all analyzed molecules. This was the case for stimulation with 10 ng/ml of IFN-γ, but also a low IFN-γ concentration of 0.1 ng/ml led to a marked upregulation of most markers. The isolated *T. cruzi* infection did not have a large effect on the phenotype of primary keratinocytes with respect to the activating ligands but induced significantly inhibitory ligands.

## Discussion

Each tissue provides a unique microenvironment with a high diversity of signals affecting immune cells. These tissue-specific factors influencing innate and adaptive immune responses are not well understood. Our study aims to fill this knowledge gap by examining early stages of *T. cruzi* infection in the skin. Considering that the skin serves as the entry point for *T. cruzi,* a local immune activation orchestrates subsequent adaptive immune responses and may set the course for the outcome of infection. We specifically focus on primary skin keratinocytes, which are the predominant cell type at the site of infection. Keratinocytes constitute approximately 95% of the cell population in the epidermis, the outermost layer of the skin (15). In acute cases of Chagas disease, patients often display a swelling in the skin, known as Romana’s sign, which results from a local inflammation at the parasite’s entry point. This swelling commonly affects the eyelid and may extend to the conjunctiva and nearby tissues. Intracellular amastigotes and immune cell infiltration can be observed in these areas. Although the Romana’s sign is a specific clinical manifestation of an acute infection with *T. cruzi*, a detailed characterization of the immunological infiltrate or a description of the early events of immune defense are lacking. Interestingly, *Ward et al*. have shown that *T. cruzi* is present in the skin of chronically infected mice, suggesting that keratinocytes might act as a reservoir for the parasite in individuals with chronic infections (3). Importantly, in immune compromised patients a chronic infection is often accompanied by a reactivation with an early skin manifestation (3, 16-19). This indicates that the skin acts a reservoir and a continuous immune surveillance controls the infection.

Our data confirms that *T. cruzi* can effectively infect primary human keratinocytes. Keratinocytes are of special interest since they are likely to be the first cell type infected upon vectorial transmission. Replication within keratinocytes and recruitment of phagocytic cells as potential vehicles for dissemination will spark the initial infection. Therefore, any immune pathway capable of lowering the initial parasitic load might have a profound effect on the subsequent course of infection. NK cells have been shown to play a crucial role in controlling *T. cruzi* infection, particularly during the acute phase. Depletion of NK cells in *T. cruzi* infected mice leads to increased susceptibility, higher parasitemia, and mortality rates (9, 20). The production of IFN-γ by NK cells is considered a major protective mechanism against *T. cruzi* infection (20–22). NK cell-mediated killing of parasites can occur through the release of granzymes, granulysin and perforin (7). Furthermore, it is crucial to note that early IFN-γ secretion orchestrates a type I immune response that activates macrophages (23). To address the interaction with *T. cruzi* infected hPEK an autologous model was employed, i.e., primary keratinocytes and NK cells were from the same donor. Degranulation was assessed via staining of CD107a on the surface of NK cells. CD107a is associated with the membranes of intracellular granules and, therefore, is present only on the cell surface following membrane fusion after degranulation (24). Upon co-incubation of NK cells with infected hPEK we observed a strong increase of surface CD107a, which was not observed when NK cells were co-incubated with non-infected hPEK. Furthermore, CD16 on NK cells was significantly downregulated after co-culture with infected hPEK compared to non-infected hPEK. A loss of CD16 occurs following activation of NK cells (25). A strong activation of NK cells is further corroborated by an increase of pro-inflammatory cytokines and cytolytic mediators in the supernatants of co-cultures. Taken together, our data provide clear evidence that NK cells are able to sense and react to *T. cruzi* infected keratinocytes. The release of granzymes, granulysin, and perforin are capable to mediate parasite killing (26), and the secretion of IFN-γ might promote a protective type I immune response and enhances the killing activity of macrophages (27, 28). To assess whether indeed NK cells exert a direct cytotoxic effect to infected hPEK the high-throughput confocal imaging system Opera Phenix® was used. This analysis revealed a reduction in the total number of cells, the infection rate, and the number of *T. cruzi* parasites per infected cell with increasing E:T ratios after co-incubation with NK cells.

A reduction in the number of infected cells might result from killing by the release of cytolytic mediators. However, it is possible that direct killing is not the primary effector mechanism employed by NK cells in *T. cruzi* infections. *Prajeeth et al*. showed that the control of *Leishmania* infections by NK cells was mediated by cytokine secretion, but not by cytotoxicity (29). In this study it was demonstrated that NK cells were not able to kill infected macrophages directly, but cytokines released by NK cells increased the antimicrobial activity of infected macrophages, such as the production of nitric oxide. Similarly, *Lieke et al.* showed that NK cells exhibited a trypanocidal effect on *T. cruzi* infected fibroblasts, which was mediated by soluble factors and by the production of nitric oxide.

Keratinocytes are known for their diverse immunological functions, including cytokine production, antigen presentation, and activation in response to pathogen invasion, cell damage, or a pro-inflammatory environment (30). To further understand their role in *T. cruzi* infection, we examined the induction of immune regulatory surface molecules known to be important for the interaction with T cells and NK cells. First, we investigated whether human primary keratinocytes undergo phenotypical changes upon infection with *T. cruzi*. In addition, keratinocytes were stimulated with IFN-γ, since this cytokine is produced by CD8^+^ T cells and NK cells in *T. cruzi* infection. It has been shown to have a protective role in acute infection but also remained elevated during the chronic stage (13, 21, 31). *T. cruzi* infection alone did not have a large effect on the phenotype of primary keratinocytes. This is surprising, since *T. cruzi* is able to trigger PRRs, and keratinocytes readily react to PRR stimulation with a pro-inflammatory immune response (30, 32). The results seen here would support the hypothesis that *T. cruzi* can enter the host relatively unnoticed. In contrast, IFN-γ stimulation induced significant upregulation of almost all analyzed surface molecules. This was the case for stimulation with 10 ng/ml of IFN-γ, but also low IFN-γ concentrations of 0.1 ng/ml led to a marked upregulation of most markers. This concentration falls within the range observed in the supernatants of NK cell co-culture experiments in reaction to *T. cruzi* infected keratinocytes (0.06 – 0.9 ng/ml). This information is presented as a fold change in Figure 3). It is important to note, however, that the actual local IFN-γ concentrations in an *in vivo* setting are difficult to assess, and the cytokine concentration is subject to degradation, uptake, and receptor binding by target cells. As the infection itself had only little effects on the surface molecule expression, it is assumed that the major phenotypical changes occurring during infection are due to IFN-γ production in the pro-inflammatory setting. Taking these experiments as basis, the infection alone would go almost unnoticed regarding the expression of HLA-ABC, HLA-DR, or ICAM-1. However, IFN-γ, but not the *T. cruzi* infection *per se*, induce an upregulation of HLA-ABC and HLA-DR. Therefore, IFN-γ stimulated keratinocytes might act as antigen-presenting cells for CD8^+^ and CD4^+^ T cells. A similar scenario is also described in previous studies (33, 34). We also observe an upregulation of ICAM-1, an adhesion molecule that is upregulated at sites of local inflammation that mediates the recruitment of leukocytes and has a direct stimulatory effect on NK cells and CD8^+^ T cells (35, 36). Black *et al.* have demonstrated an upregulation of ICAM-1 expression by keratinocytes through IFN-γ stimulation, which is in line with our results (33). Importantly, MIC-A/B was upregulated in infected as well as IFN-γ stimulated cells. MIC-A/B is an MHC class I related protein and stimulates CD8^+^ T cells and NK cells via the receptor NKG2D (37).

Of particular interest are the expression of the inhibitory receptors PD-L1 and PD-L2, since both can inhibit immune cell activation by binding to PD-1, which is expressed on T and NK cells (38–40). Both, PD-L1 and PD-L2 are upregulated in *T. cruzi* infected and IFN-γ stimulated keratinocytes. While an interaction of PD-L1 and/or PD-L2 with PD-1 is typically associated with a reduced immune response, it might also be a counter regulatory mechanism for limiting tissue damage in an infectious setting (41). This is in accordance with the finding that a blockade of PD-1/PD-L1 interaction was associated with increased heart pathology in murine models of acute and chronic *T. cruzi* infection (42, 43). On the other hand, PD-L1/-L2 expression could impede parasite clearance by preventing an effective immune response. For instance, Gutierrez et al. observed a reduced parasite load and augmented inflammatory response upon PD-1/PD-L1 blockade (43).

HVEM is another receptor that was induced upon IFN-γ stimulation. Depending on the ligand, HVEM can mediate either activating or inhibitory signals. Ligation to CD160 on T cells, for example, has an inhibitory effect, whereas binding to CD160 on a NK cell leads to their activation (44, 45). An increased surface expression of CD160 on CD4^+^ and CD8^+^ T cells has been found in acutely and chronically *T. cruzi* infected mice (46). Other inhibitory receptors expressed by CD8^+^ T cells and NK cells are Tim-3 and TIGIT. Their respective ligands, Galectin-9 and CD155, were found to be upregulated by keratinocytes upon IFN-γ stimulation in this study. To our knowledge, induction of these molecules on keratinocytes has not been addressed by any studies so far. It is intriguing to speculate that these might serve as potential counter-regulatory mechanism to prevent inflammation of the skin.

Since we observed the induction of inhibitory and activating ligands, it is not clear which effect dominates. However, since we observed a strong activation of NK cells and an induction of their antimicrobial activity to infected keratinocytes at least for NK cells activating pathways are dominating. This may have the potential to strongly reduce the parasite inoculum during the initial infection, which might have a profound effect on the later course of infection.

## Materials and Methods

### Isolation of Peripheral Blood Mononuclear Cells

Blood was drawn from the median cubital vein using Lithium-Heparin tubes and stored for a maximum of 8h at RT. Peripheral Blood Mononuclear Cells (PBMCs) were isolated from whole blood of healthy volunteers using SepMate™ tubes as described by *Mackroth et al.* (47). PBMCs were cryopreserved or used for co-cultivation experiments.

### Isolation of Primary Keratinocytes

Keratinocytes were isolated from human 6 mm biopsies kindly provided by healthy donors. Obtaining ethics approval was not required. Donors gave informed consent before the donation of skin and matching blood samples. The age and sex of the donors, as well as the biopsy site, are listed in Table 1. Biopsies were stored at 4 °C in a physiologic solution for a maximum of 48 h. Under sterile conditions, primary keratinocytes were isolated as described in (48). Briefly, hair and fatty tissue were removed, and tissue was washed with PBS, rinsed in 70 % ethanol for 10 sec, and washed again in PBS. The skin was cut into 2-3 mm thin stripes and incubated overnight at 4 °C in Dispase II. On the next day, the epidermis was separated from the dermis and collected in PBS. Single cells were obtained by incubating the epidermis in 1 ml of trypsin-EDTA solution for 5 min at 37 °C, followed by mechanical segregation. Digestion was stopped and after 5 min centrifugation at 200 g, the supernatant was aspirated and cells were resuspended in keratinocyte growth medium (KGM). Cells were cultivated in T25 cell flasks or stored in cryovials in liquid nitrogen at −196 °C.

**Table 1.** Information about the donors.

| Primary Keratinocytes ID | Age (years) | Sex | Biopsy site |
| --- | --- | --- | --- |
| PK1 | 39 | female | upper arm |
| PK2 | 54 | male | upper arm |
| PK6 | 55 | female | inguinal region |

### Cultivation of Keratinocytes

Keratinocytes were cultivated in keratinocyte growth medium (KGM; DermaLife K medium complete kit (Lifeline Cell Technology), 100 U/ml Penicillin, 100 U/ml Streptomycin (Capricorn Scientific) in T25 cell culture flasks at a density of 0.2 - 0.5 x 10^6^ cells per flask or in T75 cell culture flasks at a density of 0.5 −1.10^6^ cells per flask. For cultivation, 1 µg/ml of phenol red (Sigma-Aldrich) was added to the medium. The flasks were incubated at 37 °C and 5 % CO_2_ atmosphere. Cells were passaged at a confluency of > 80 % and the medium was changed in case confluency was not reached after 3-4 days. Primary keratinocytes were passaged by washing the cell layer with PBS, followed by 10-15 min incubation with Trypsin-0.1 % EDTA (Capricorn Scientific) at 37 °C. The keratinocytes were used for assays at passages 3-4. Keratinocytes were tested negative for mycoplasma.

### Cryopreservation

Keratinocytes were cryopreserved by resuspending 0.5 - 1 x 10^6^ cells per 1 ml of freezing medium. PBMCs were frozen at 1 x 10^7^ cells per 1 ml of freezing medium (FCS, 10 % DMSO (Sigma-Aldrich)). Keratinocytes were thawed by resuspension in pre-warmed washing medium (DMEM (PAN Biotech), 10 % FCS, 100 U/ml Penicillin, 100 U/ml Streptomycin, 2 mM L-Glutamine (Capricorn Scientific)).

#### Parasite Culture

All work involving viable *T. cruzi* parasites was conducted under biosafety level 3 conditions (BSL-3). Glioblastoma 86HG39 cells were used to keep parasites alive *in vitro* and *T. cruzi* parasites were passages as described in (13).

#### Infection of Keratinocytes

Keratinocytes were seeded one day before infection. The number of cells that were seeded per well were optimized for confluency of approx. 60 % 24 h after seeding. Additional wells were seeded per plate for counting of cells before infection, as well as before the addition of NK cells. Before *T. cruzi* parasites were added, the medium was changed. Non-infected controls were included in all experiments.

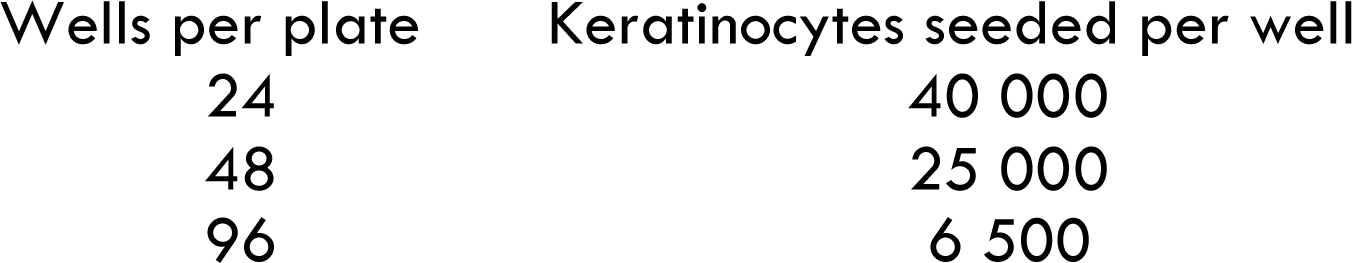

The infection with *T. cruzi* parasites was performed as described in (13). The parasites were resuspended in an appropriate volume of pre-warmed KGM. The MOI of 1:1, 3:1, or 6:1 was used to determine the infection rates of *T. cruzi* Brazil and *T. cruzi* Tulahuen in primary keratinocytes. For all other experiments, an MOI of 3:1 was used. The keratinocytes were co-cultivated with the parasites in KGM for 24 h at 37 °C and 5 % CO_2_ to allow parasite entry. 24 h after infection, the wells were washed once with medium to remove free parasites, and fresh medium was added.

### Immunofluorescence Staining

Keratinocytes were seeded on chamber slides and infected with *T. cruzi* 24 h later. Every 24 h up to 96 h post-infection (p.i.), chamber slides were washed twice with PBS and the cells fixed with 4 % PFA for 45 min at RT. The chamber slides were stored in 200 µl of PBS per well at 4 °C.

For staining, the cells were permeabilized with 0.1 % Triton-X-100 in PBS for 5 min, followed by three washing steps with PBS. Cells were incubated with 100 µl of rabbit anti-*T. cruzi* serum (1:2000) and anti-cytokeratin pan antibody (1:100, Thermo Fisher Scientific) in PBS for 1.5 h at 37 °C. Following three washing steps with PBS, the cells were incubated with 100 µl of secondary antibody mix for 45 min at RT protected from light. The secondary antibody mix contained Alexa Fluor 568 conjugated goat anti-rabbit IgG antibody (1:200) and Alexa Fluor 488 conjugated goat anti-mouse IgG antibody (1:200, both from LifeTechnologies) in PBS. The wells were washed PBS, after which the chambers were removed from the slide. One drop of Roti®-Mount FluorCare DAPI (Carl Roth) was added per well and a coverslip was applied. The slides were imaged using a Zeiss Axio Imager M.1 fluorescence microscope. Image overlays were created using the FIJI ImageJ software.

### Immunofluorescence Staining for High Content Screening

The High Content Screening (HCS) system Opera Phenix® and Harmony® software(version 4.6 from PerkinElmer) were used to determine the total number of cells, the infection rate and the number of trypanosomes in the infected keratinocytes. Cells were seeded in black 96-well CellCarrier™ plates and infected with *T. cruzi* Brazil or *T. cruzi* Tulahuen 24 h later. At different time points, the wells were carefully washed twice with pre-warmed PBS and fixed for 45 min with 4 % PFA at RT for analysis. After washing with PBS, the plates were stored at 4 °C in 200 µl of PBS. For immunofluorescence staining, the wells were washed twice with HCS wash buffer (0.1 % Triton-X-100 (Sigma-Aldrich) in PBS) for 5 min. All washing and incubation steps were performed under gentle shaking at 300 rpm and RT. The cells were then permeabilized with HCS permeabilization buffer (PBS, 0.1 % Triton-X-100, 50 mM NH_4_Cl (Carl Roth)) for 15 min, followed by 30 min blocking with HCS blocking solution (PBS, 0.1 % Triton X-100, 2 % BSA (SERVA Electrophoresis)). *T. cruzi* was stained by adding mouse anti-*T. cruzi* serum diluted 1:2000 in blocking solution for 90 min. After three washing steps with wash buffer for 5 min each, 60 µl of the secondary antibody mixture were added and incubated for 60 min protected from light. As a secondary antibody, anti-mouse IgG AF647 (Life Technologies) was used at 1:8000 dilution in blocking solution together with 0.01 mg/ml DAPI (Thermo Fisher). The wells were washed with wash buffer and PBS. The plate was stored in 200 µl of PBS at 4 °C protected from light until it was measured at the Opera Phenix®.

The Opera Phenix® is a confocal imaging system that automatically creates several images per well. Each well is divided into an 11×11 grid, from which 15 sections were selected for image acquisition. The images can be analyzed with the Harmony® software and a custom made image analysis sequence. Here, the output generated by the analysis sequence included the number of cells, percentage of infected cells, number of trypanosomes, and trypanosomes per infected cell. Cells were detected via the nucleus and a weak cytoplasmic background signal that originates from the unspecific binding of the primary antibody. The *T. cruzi* parasites were detected via a strong AF647 signal together with a small DAPI stained spot reflecting the parasitic nucleus. Several parameters were included to reliably detect *T. cruzi* parasites and the full analysis sequence can be found in Table S3.

### Stimulation of Primary Keratinocytes

Primary Keratinocytes from three donors were seeded in 24-well plates and stimulated with KGM Media supplemented either with 0.1 ng/ml or 10 ng/ml of recombinant human IFN-γ (PeproTech), or infected with *T. cruzi* Brazil. The keratinocytes were stained for various surface markers using flow cytometry. The MOI used was 3:. Furthermore, if cell counts allowed it negative control per donor were included as neither infected nor stimulated. The plates were incubated for 72 h at 37 °C and 5 % CO_2_. 24 h p.i., the medium from all wells was removed, the wells were washed once with KGM, and fresh KGM was added. 72 h p.i., the keratinocytes were washed with cold PBS and detached by 15 min incubation at 37 °C with 100 µl/well of trypsin (TrypLE™, Life Technologies). The reaction was stopped by adding 150 µl of DMEM supplemented with 10 % FCS, after which the cells were thoroughly resuspended and transferred to a 96-well U-bottom plate for flow cytometry staining.

### Isolation of NK cells and Co-Culture Experiments

NK cells were isolated from frozen PBMCs by negative selection using the MojoSort™ Human NK Cell Isolation Kit (BioLegend) following the manufacturer’s instructions. The NK cells were counted and were resuspended in cRPMI supplemented with 5 ng/ml rhIL-15 (PeproTech) and rested overnight in a 96-well U-bottom plate at 37 °C and 5 % CO_2_.

Subsequently, NK cells were incubated with *T. cruzi* Brazil infected and non-infected primary keratinocytes. The keratinocytes were washed once with cRPMI, and the NK cells were added to cRMPI supplemented with 1 ng/ml rhIL-15. For flow cytometric analysis, the NK cells were added at an E: T ratio of 1:10. For supernatant and HCS analysis, E:T ratios of 1:3 and 10:1 were used, respectively. After the NK cells were added, the plates were centrifuged at 100 x g for 30 sec to accelerate the contact between the target and effector cells. The NK cells were co-cultured with the keratinocytes for 24 h. For the last 5 h of co-culture, Brefeldin A and APC-Cy7 conjugated anti-CD107a antibody (1:200) were added to the wells for flow cytometry staining. The plate was centrifuged for 30 sec at 100 x g to re-establish cell contact. After 24 h of co-culture, the NK cells were harvested and analyzed using flow cytometry. The co-culture supernatants were collected and analyzed using a multiplex bead assay, Briefly, the NK cells in the 24-well plate were harvested by pipetting the culture supernatant up and down and transferring it to a 96-well U-bottom plate for flow cytometry stainingWednesday, August 16, 2023. For cell culture supernatants, plate was centrifuged for 5 min at 500 x g and 4 °C, after which the supernatants were carefully transferred to a 96-well U-bottom plate. The plate was centrifuged for 15 min at 2671 x g and 4 °C to pelletize any parasites, and the supernatant was stored at −20 °C. For HCS analysis, the CellCarrier™ plate was washed and fixed as described above.

### Flow Cytometry Staining

The flow cytometry staining was performed as described in (13). Briefly, all washing steps were performed at 4 °C for 5 min at 400 x g (keratinocytes) or 500 x g (NK cells). For each staining, a negative control, as well as an unstained control were performed. First, a LIVE/DEAD™ fixable blue stain was performed. Subsequently, cells were washed with FACS buffer (PBS, 2 % FCS, 2 mM EDTA), surface staining antibodies were added and incubated for 30 minutes at 4°C. For intracellular cytokine staining, cells were fixed and permeabilized using the Foxp3/Transcription Factor Buffer Set (ThermoFisher) for 45 minutes at RT. Following fixation and permeabilization, intracellular staining was performed for 30 minutes at 4°C. Measurements were taken using Cytek® Aurora spectral flow cytometer and SpectroFlo® software. The data was analyzed using FlowJo™. Gates were established based on fluorescence minus one (FMO) controls. The gating strategy is depicted in S3 and S6. Table 2 provides a list of antibodies used in flow cytometric analyses.

**Table 2.**
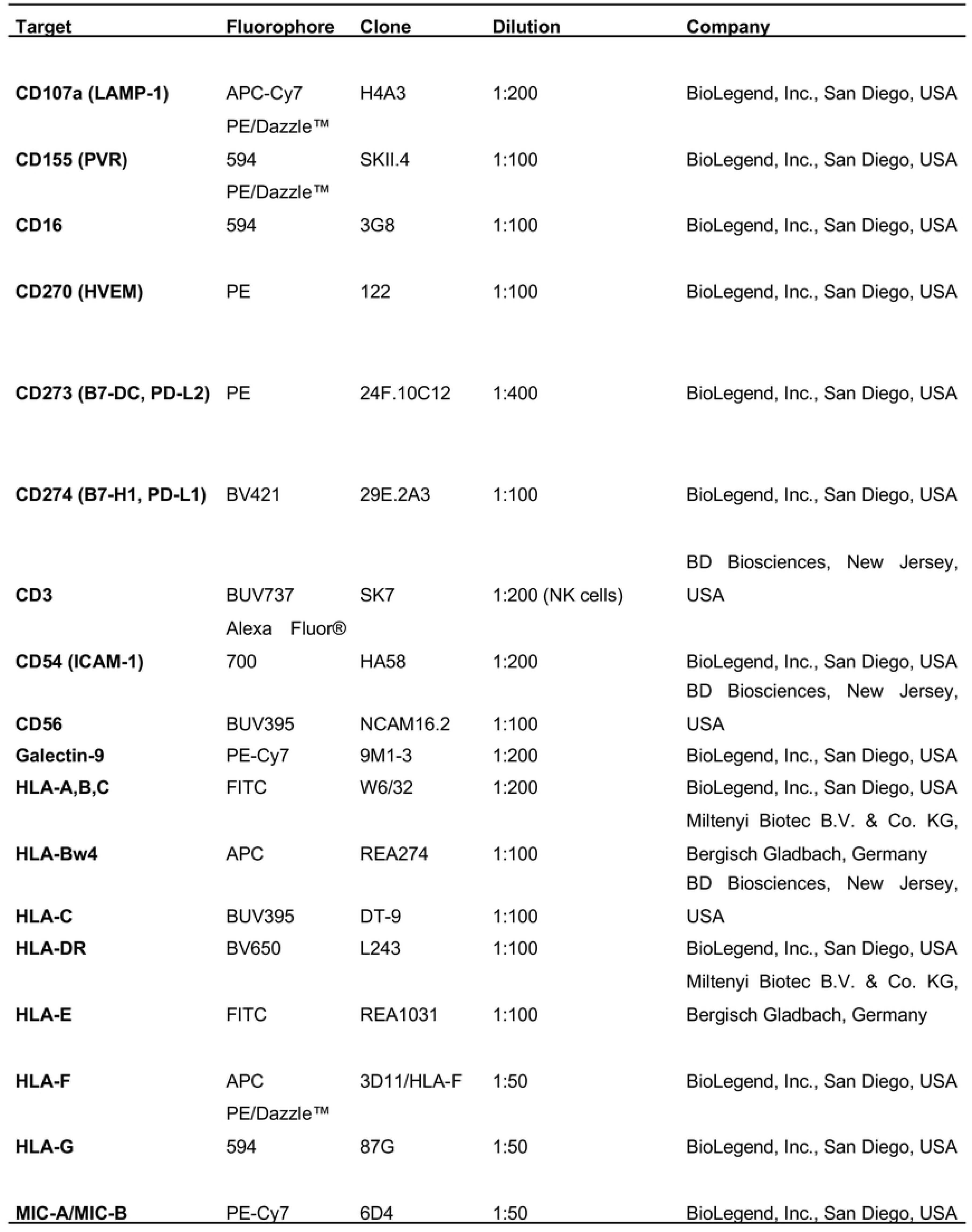
List of Antibodies.

**Table 3.**
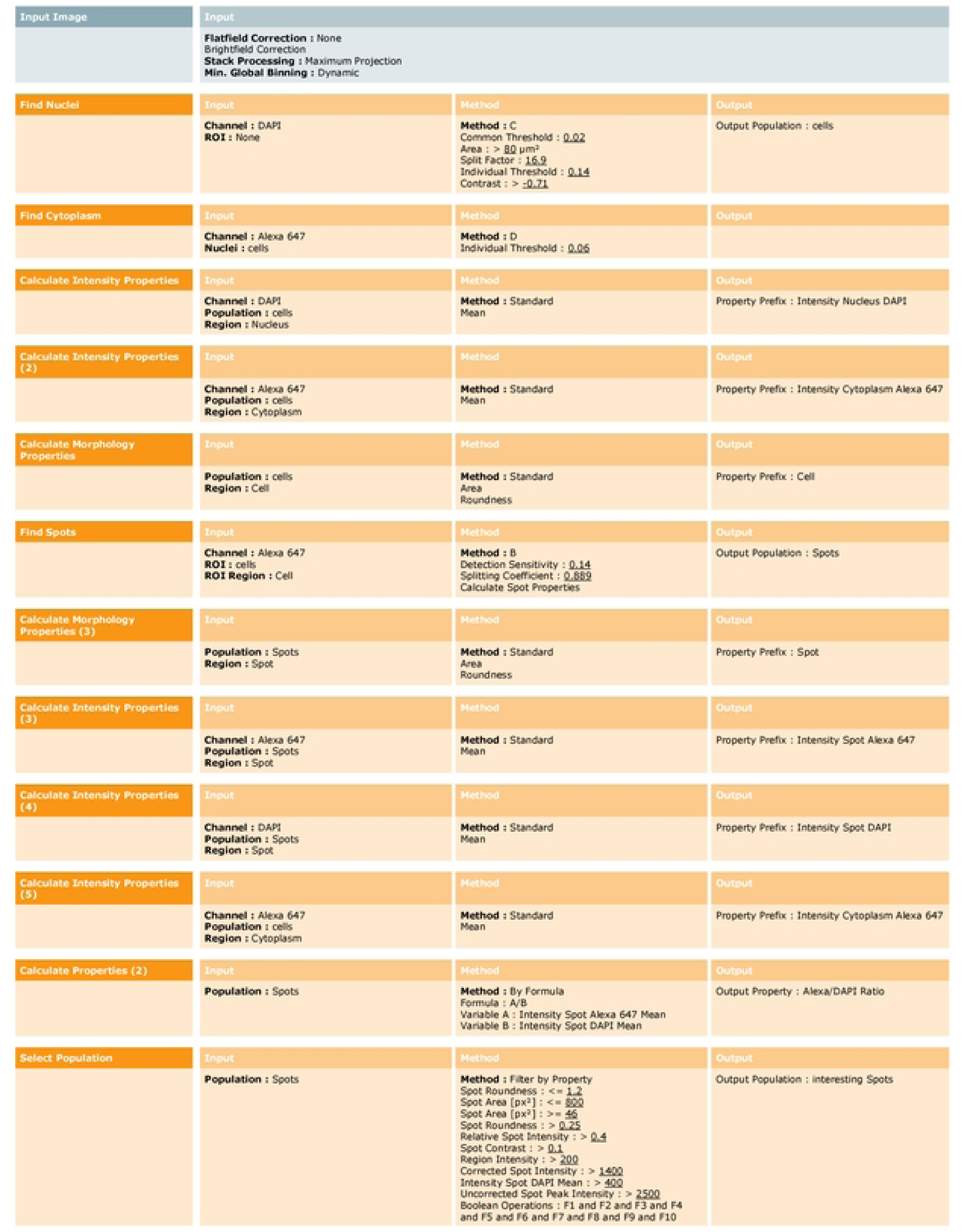
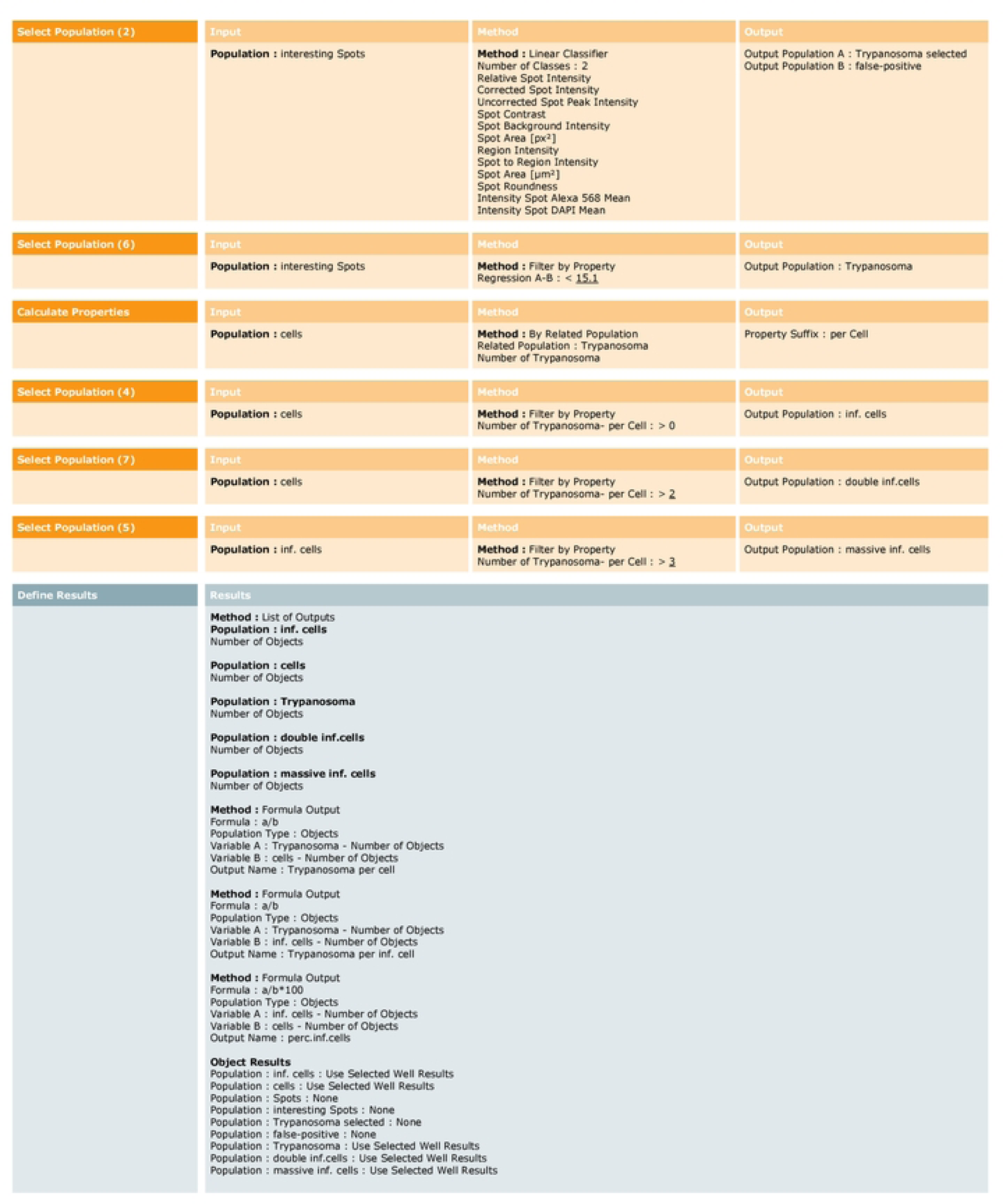
The customed analytical sequence was developed employing Harmony® software version 4.6 and can be employed for images produced using the Opera Phenix® confocal imaging system.

### Cytokine Measurement

Cytokines were quantified using the LEGENDplex Human CD8/NK Panel 13-plex (Cat.No # 740267 Lot B330268) from Biolegend. This is a bead-based immunoassay able to quantify the concentration of different analytes in co-culture supernatants. The assay was performed according to the manufacturer’s instructions and measured using at the BD Accuri™ C6 flow cytometer. The data was analyzed using the LEGENDPlex™ Data Analysis Software Suite.

### Statistics

Statistical analysis was performed using GraphPad Prism 9. The data were tested for normal distribution using a Shapiro-Wilk normality test. An unpaired t-test with Welch’s correction was performed to evaluate the differences between the two groups. In case the data did not pass the normality test, a Mann-Whitney test was performed. For analysis of more than two groups, Brown-Forsythe and Welch ANOVA and Dunnett’s T3 multiple comparison tests were used for normally distributed values. Kruskal-Wallis and Dunn’s multiple comparison tests were used for values that did not follow a normal distribution. Results were rated as statistically significant when p <0.05.

### Conflict of Interest Statement

The authors declare that the research was conducted in the absence of any commercial or financial relationships that could be construed as a potential conflict of interest.

## Acknowledgments

We are grateful to the healthy volunteers who participated in the study. We thank Christiane Steeg for technical assistance and Dr. Gregory Williams for assistance language editing.

## Author Contributions Statement

Conceptualization: TJ & RIG; Formal Analysis & Methodology: KK, JB, HF, RIG; Funding acquisition: TJ; Resources: TJ, HL, BV, AV; Writing: KK, RIG; Supervision: TJ, RIG. All authors collaborated in the editing process and accepted the submitted version of the manuscript.

## Supplementary Figures

**Sl.**
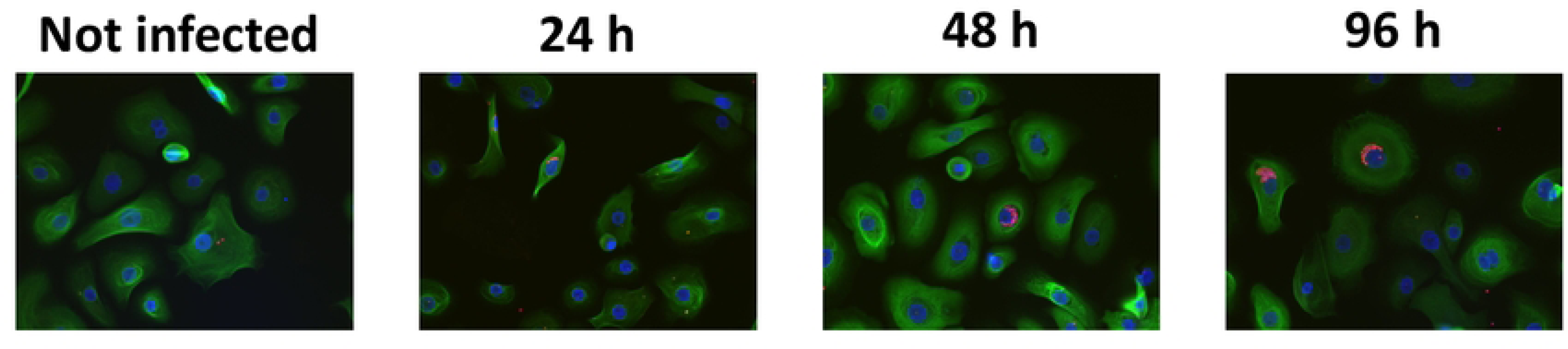
Infection of human primary keratlnocytes with *T.cruzl* Tulahuen. Primary keratinocytes were infected with *T. cruzi* Tulahuen at a MOI of 3:1 for 24 h, 48 h, and 96 h. Keratinocytes (green) and trypanosomes (red) were visualized by indirect immunofluorescenceusing a pan anti-cytokeratin antibody, polyclonal anti-T. *cruzi* serum, and DAPI. Images were obtained at 200x magnification. n.i., not infected; TcT inf.,*T. cruzi* Tulahuen infected. Primary keratinocytes were infected with *T. cruzi* Tulahuen at a MOI of 3:1 for 24 h, 48 h, 72 h, and 96 h. Keratinocytes (green) and trypanosomes (red) were visualized by indirect immunofluorescence using an pan anti-cytokeratin antibody, polyclonal anti-T. *cruzi* serum, and DAPI. Images were obtained at 200x magnification. n.i., not infected; TcT inf., *T. cruzi* Tulahuen infected.

**S2.**
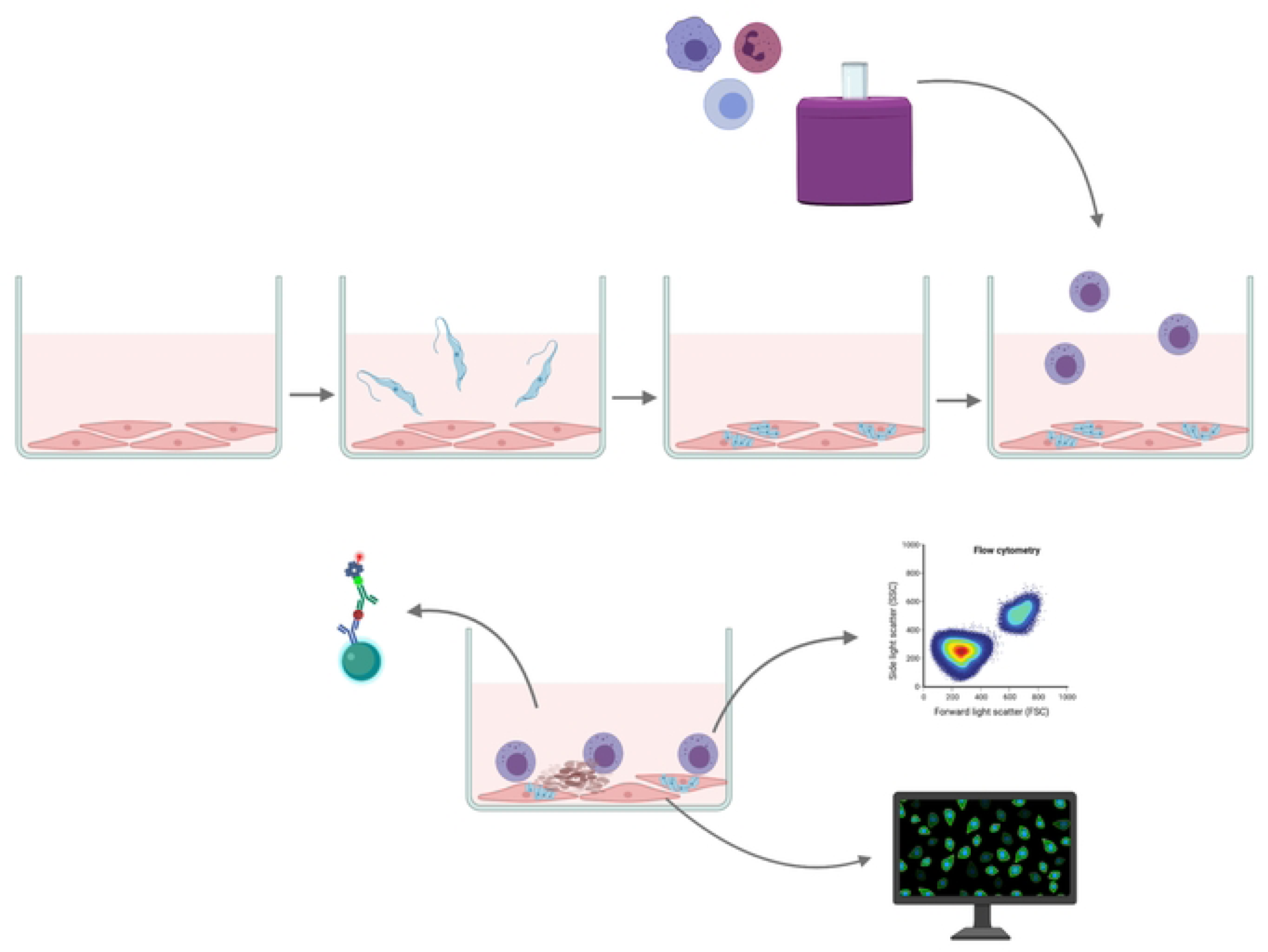
Scheme of autologous co-cultivation. Figure created with a licensed version of Biorender.

**S3.**
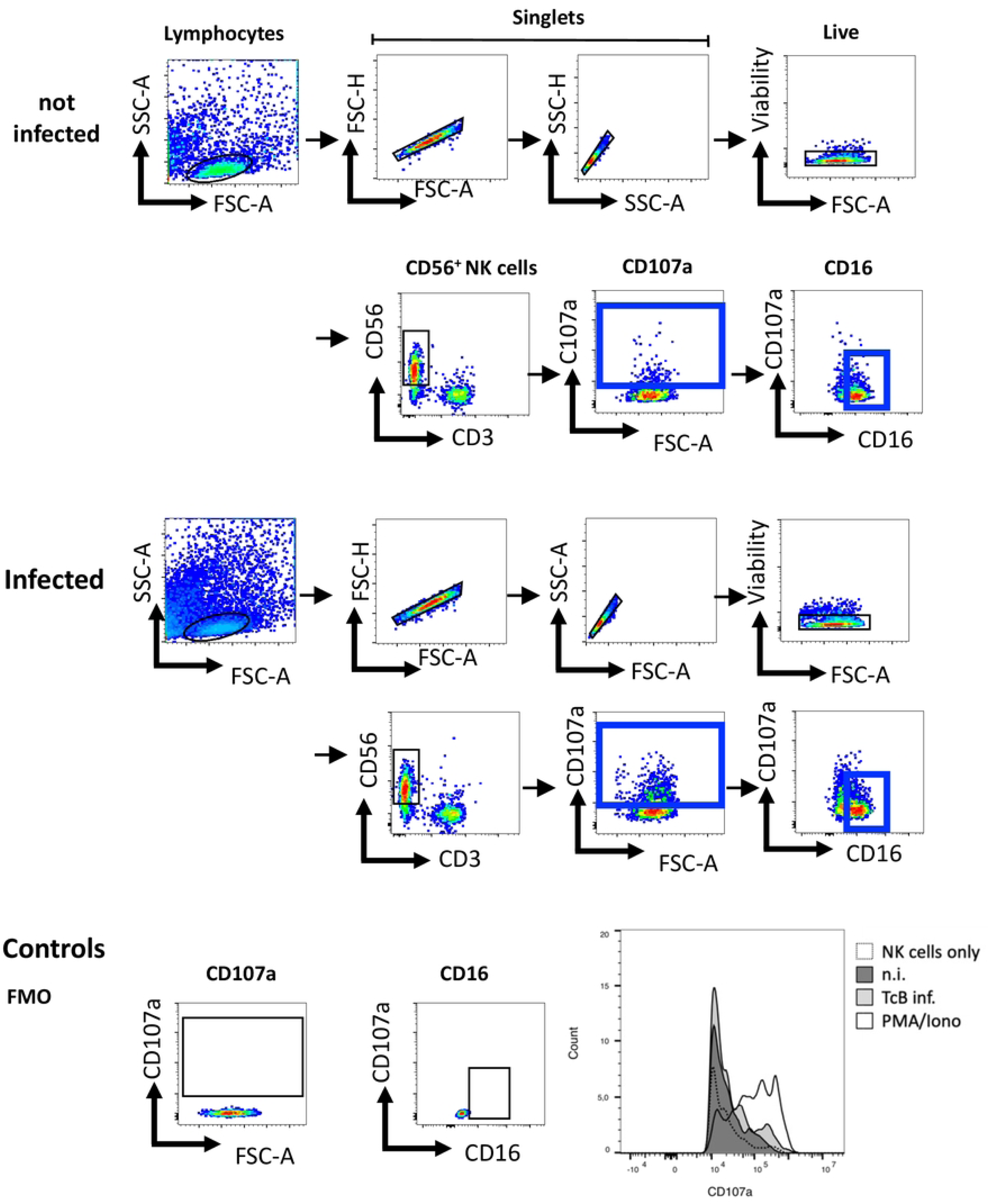
Gating strategy for NK cells. The gating strategy for analyzing NK cells after being co-cultured with autologous keratinocytes involves defining NKcells based on the expression of CD3neg and CD56+. Within the NK cell population, the gating includes CD107a and CD16 (highlighted in blue). The representative gating strategy is shown in the upper panel for co-culture with uninfected keratinocytes and in the middle panel for co-culture with infected keratinocytes. The gates are based on the respective fluorescence minus one (FMO) control, as shown in the lower panel. Additionally, the representative histograms depict the expression of CD107a.

**S4.**
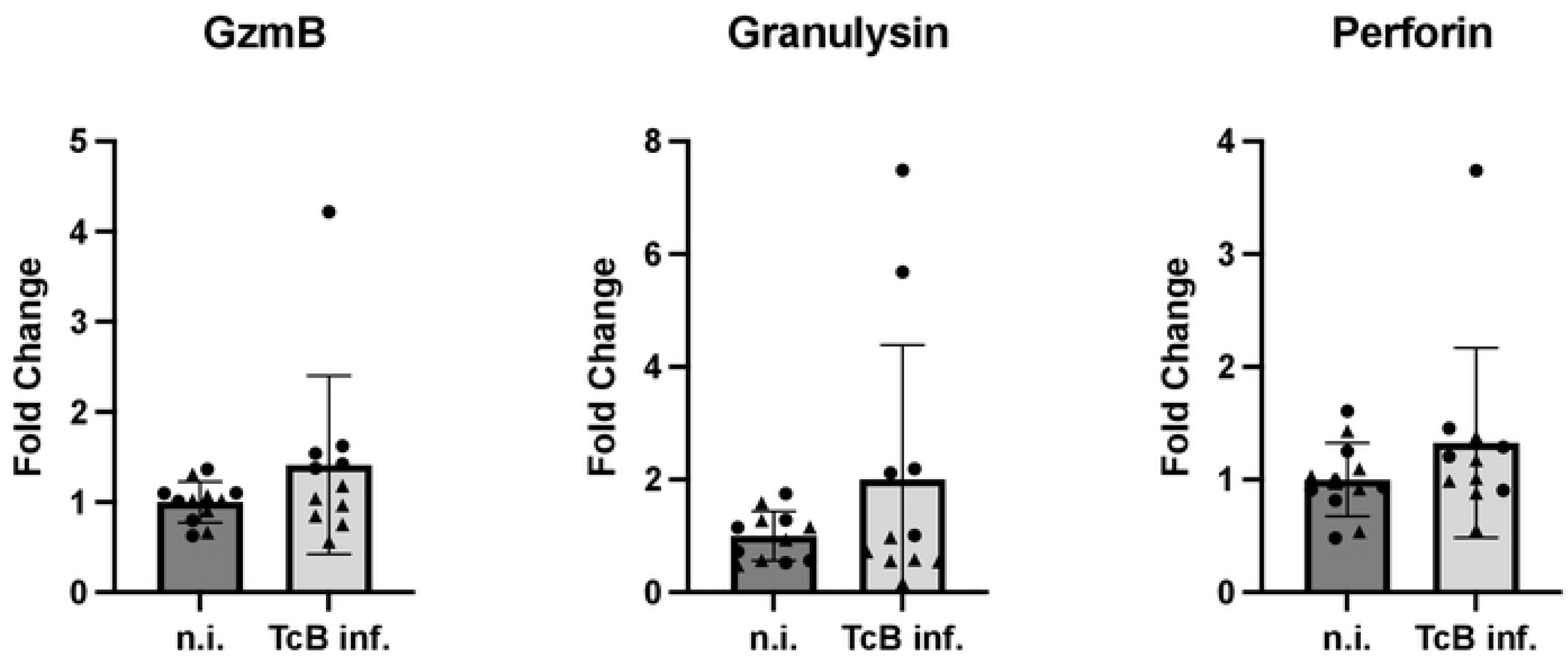
Analysis of cytoklnes and cytolytic mediators.

**SS.**
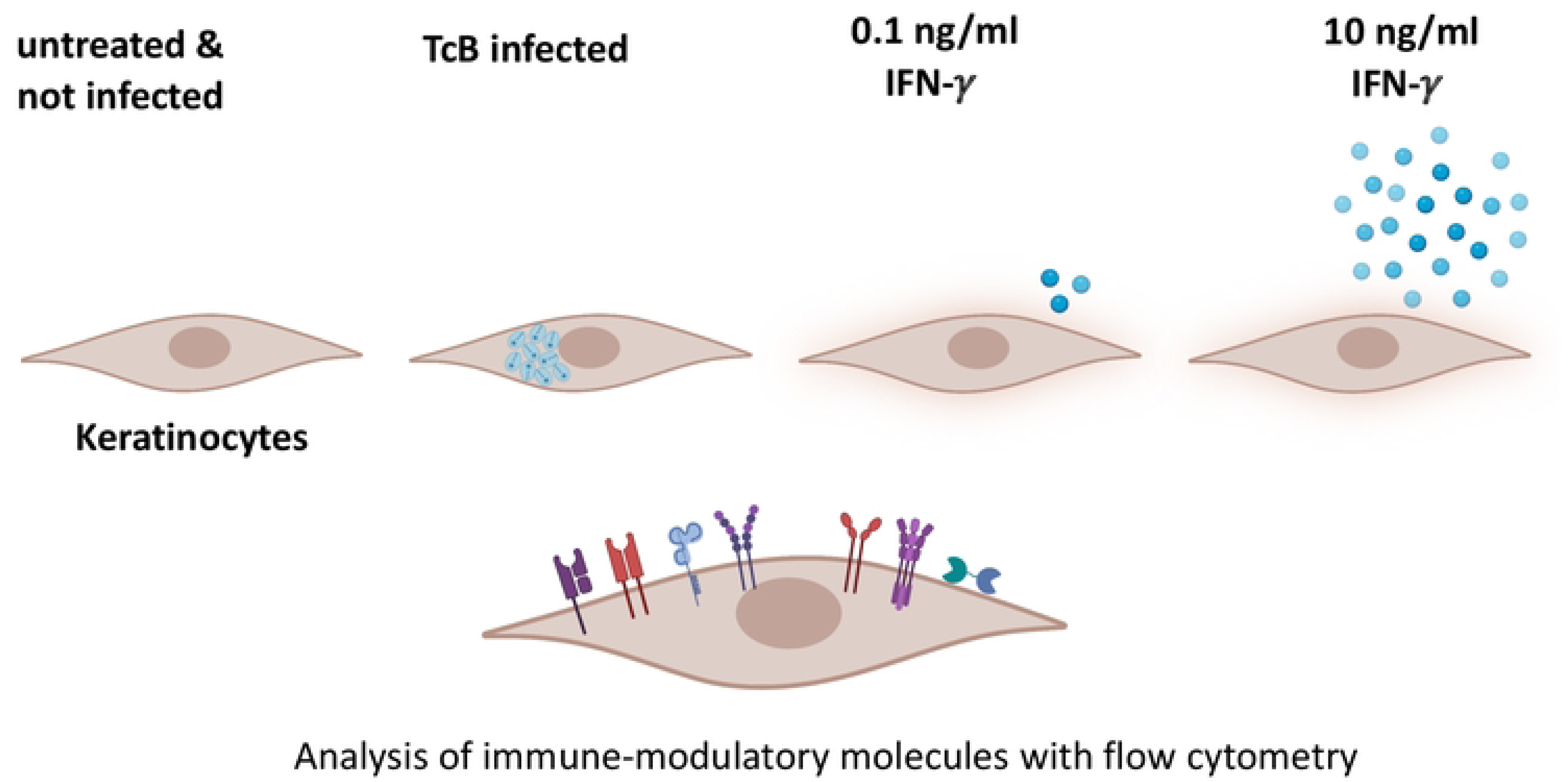
Scheme of experimental infection of human primary keratinocytes withT. *cruzi* and IFN-y stimulation.

**S6.**
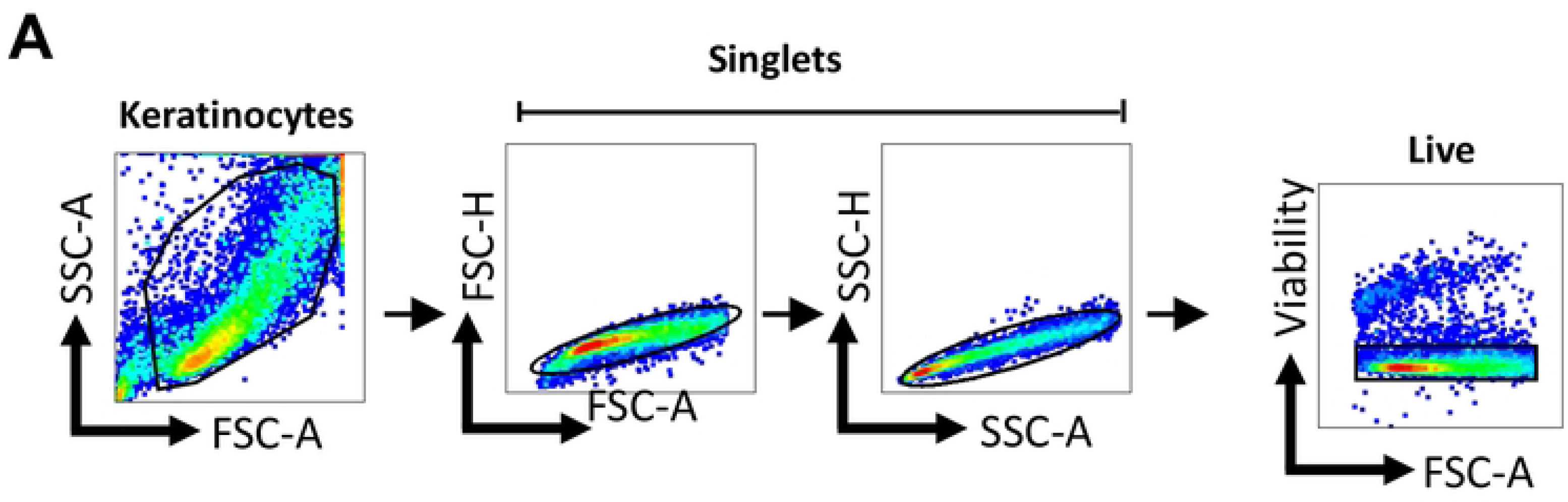
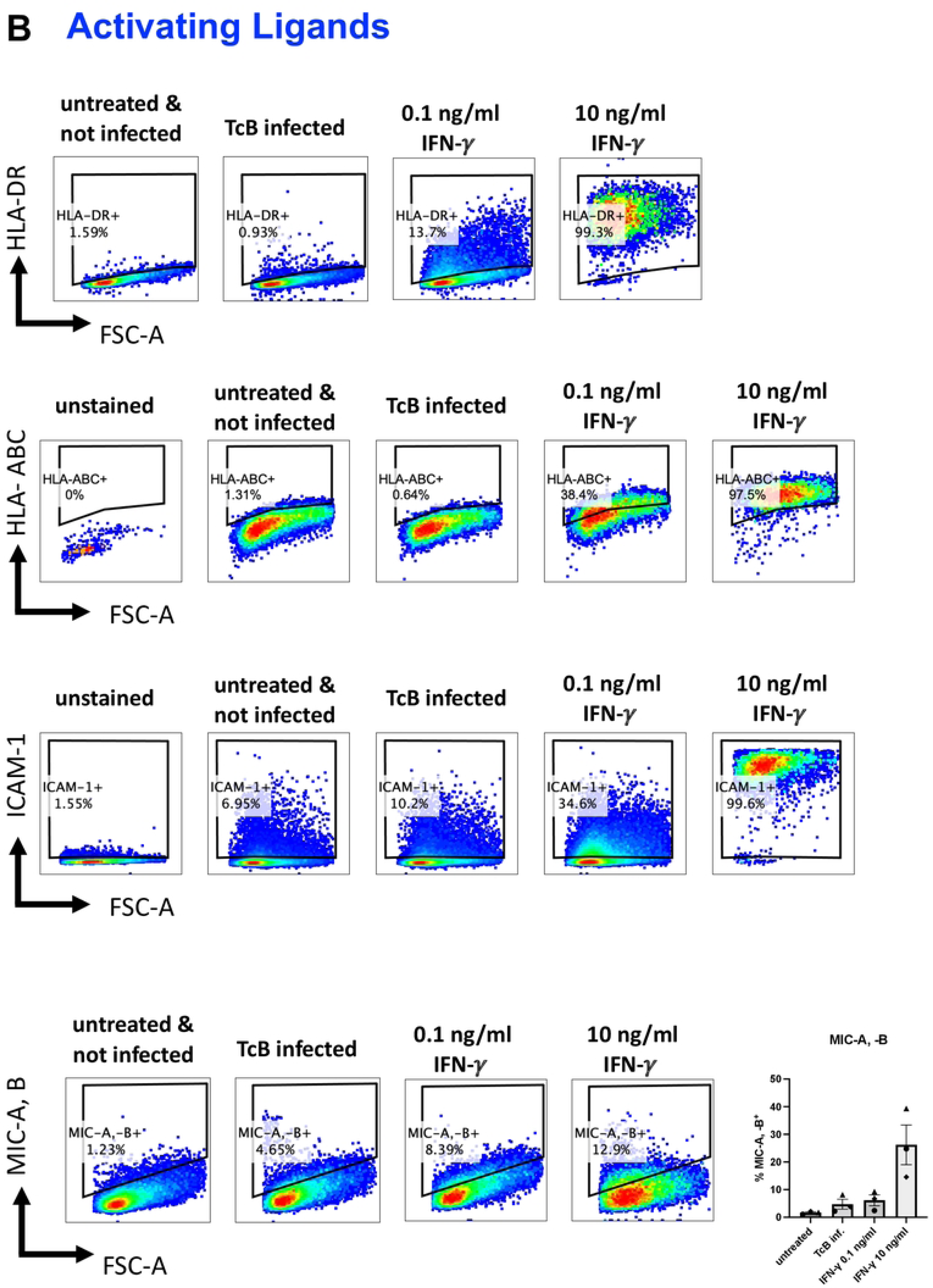
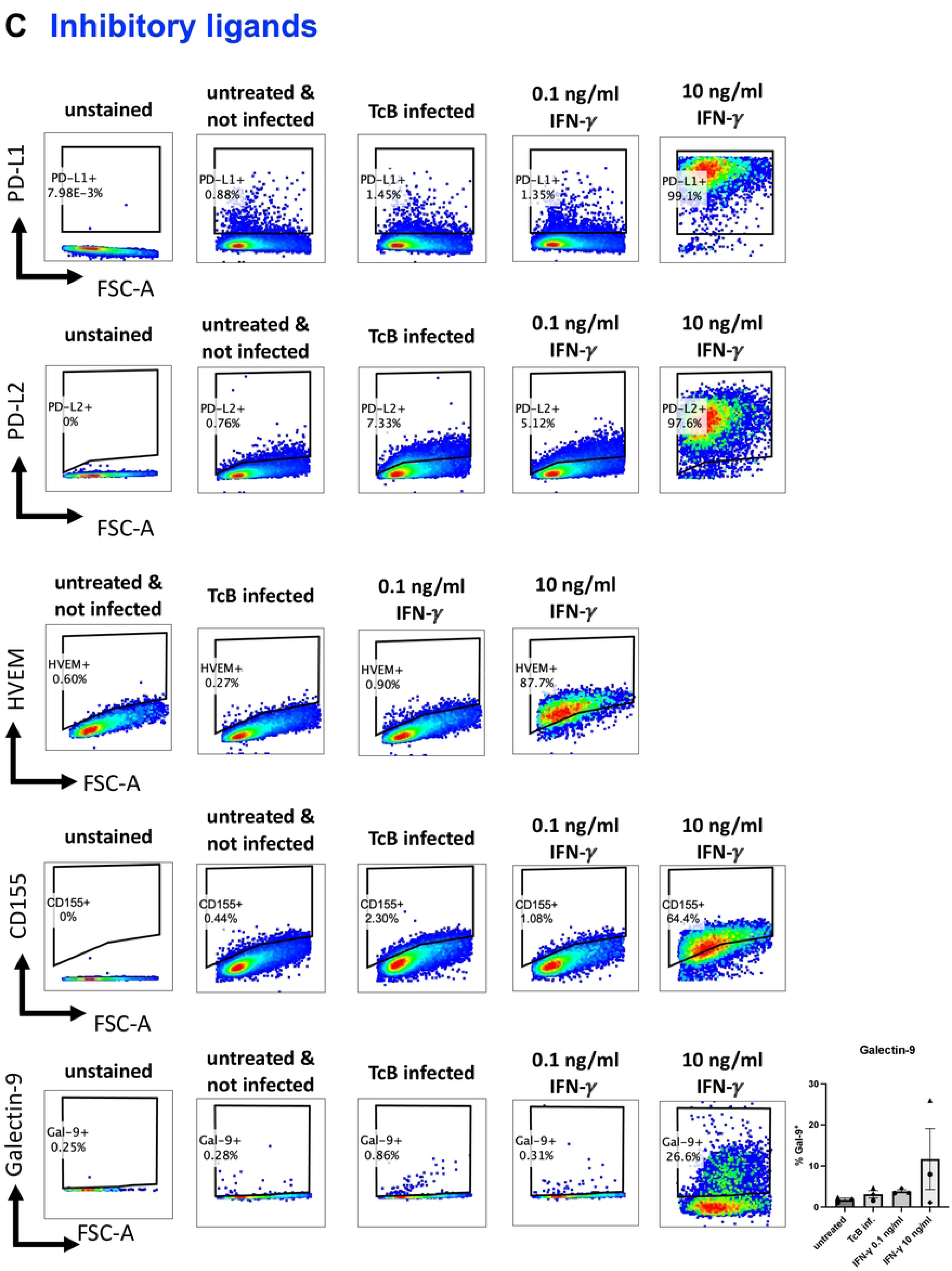
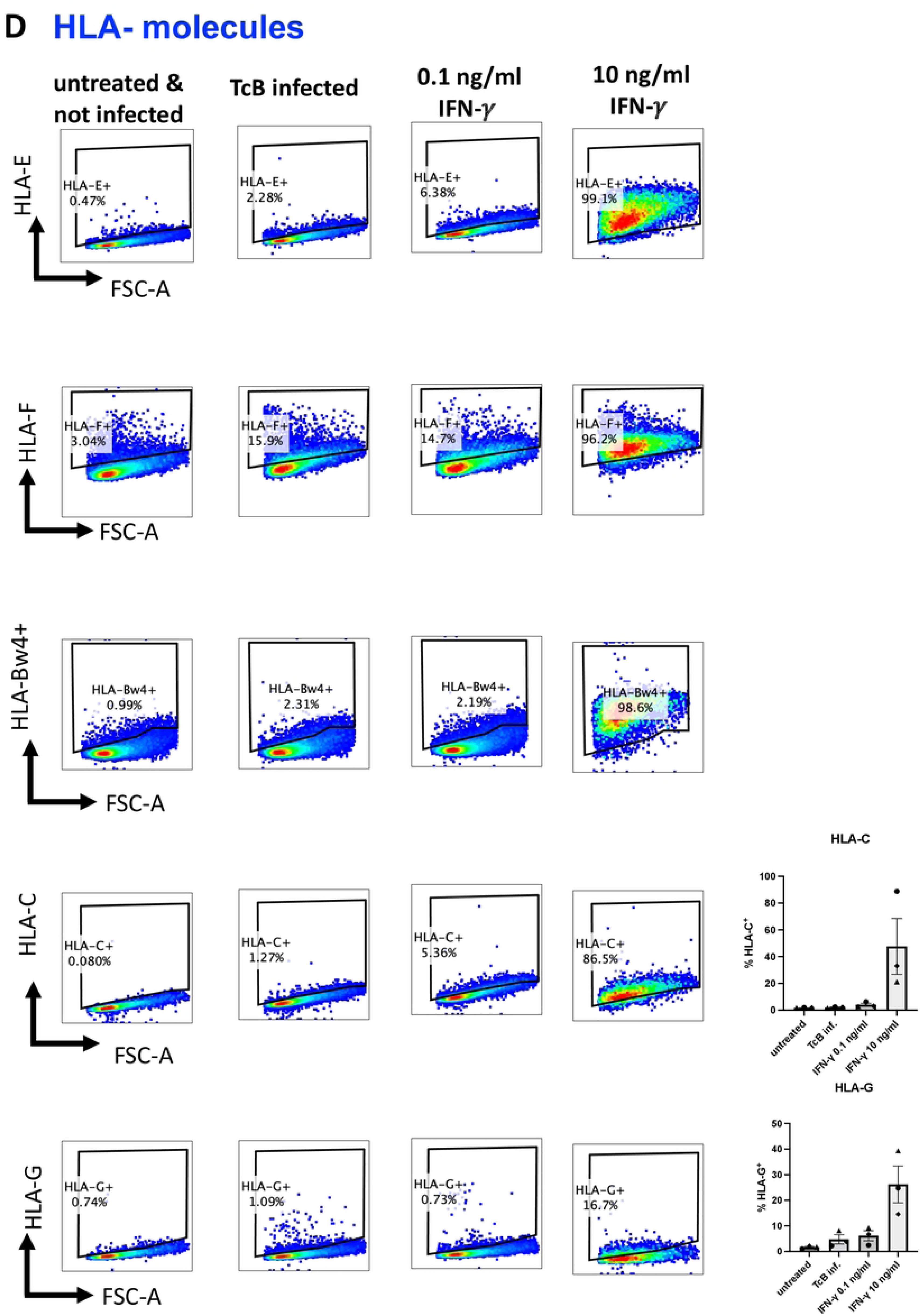
Representative gatingstrategy of flow cytometric analysis of immune modulatory molecules on the surface of keratlnocytes. A) Representative gating strategy for hPEK based on physical parameters and viability dye.All ligands have been gated on living hPEK. The plots that follow display the representative strategy of various groups, including the unstained and untreated uninfected controls, as well as the TcB infected group treated with either 0.1 ng/ml or 10 ng/ml IFN-y. B) Activating ligands HLA-DR, HLA-ABC, ICAM-1 and MIC-A, B. C) Inhibitory ligands: PD-ll, PD-l2, HVEM, CDlSS, Galectin-9. D) HLA-Molecules: HLA·E, HLA·F, HLA-Bw4+, HLA-C, HLA-G.

## Notes

### Competing Interest Statement

The authors have declared no competing interest.

